# The Wood Image Analysis and Dataset (WIAD): open-access visual analysis tools to advance the ecological data revolution

**DOI:** 10.1101/2020.12.16.423133

**Authors:** Tim Rademacher, Bijan Seyednasrollah, David J. Basler, Jian Cheng, Tessa Mandra, Elise Miller, Zuid Lin, David A. Orwig, Neil Pederson, Hanspeter Pfister, Donglai Wei, Li Yao, Andrew D. Richardson

**Author notes:** Corresponding author, 324 North Main Street, Petersham, 01366, Massachusetts, USA.

## Abstract

1. Ecological data is collected and shared at an increasingly rapid pace, but it is often shared in inconsistent and untraceable processed forms. Images of wood contain a wealth of information such as colours and textures but are most commonly reduced to ring-width measurements before they can be shared in various common file formats. Archiving digital images of wood samples in libraries, that have been developed for ecological analysis and are publicly available, remains the exception.
2. We developed the Wood Image Analysis and Dataset (WIAD) an open-source application including a web-interface to integrate basic visual analysis of wood samples, such as increment cores, thin sections or x-ray films, basic data processing, and archiving of the images and derived data to facilitate transparency and reproducibility in studies using visual characteristics of wood.
3. WIAD provides user-friendly tools to manipulate images of wood samples, mark and measure wood characteristics such as growth increments, density fluctuations, early- and latewood widths, and fire scars, and to visualise, process, and archive images, metadata, and the derived data.
4. WIAD constitutes a step towards the reproducible automation of tree-ring analysis, while establishing an open-source foundation to create improved community developed repositories which would enable novel ecological studies harnessing the wealth of existing visual data.

## 1. Introduction

Ecological data is collected at an increasingly rapid pace, but to optimise its use data needs to be shared in its raw format with accompanying ancillary and metadata (Michener, 2015). While the community of science using tree rings has been leading data sharing efforts (e.g., International Tree Ring Database and DendroEcological Network), there is still an urgent need for improved tree-ring archives (Babst et al., 2017; Zhao et al., 2018). Recently, digital archives have been developed to accommodate the need to share raw images of wood samples and meet the challenge of image size (e.g., DendroElevator). However, there are still no open-source analysis tools that easily interface with such archives. Wood samples, such as increment cores, cross sections and microcores are mostly still prepared according to standard procedures, extracting visual properties such as ring-width series (Fritts, 1976), blue intensity as a proxy for wood density (Björklund et al., 2014), or cellular dimensions (Björklund et al., 2017; von Arx & Carrer, 2014). The rich information embedded in the colours, textures and structures of each wood sample, which are used to identify these visual features in the first place, are generally collapsed to a simple one-dimensional geometric measurement (i.e., ring width) before they are shared in one of many common file formats (Brewer et al., 2011).

While multiple specialised tools for digital analysis of wood images exist (e.g., WinDendro, ROXAS, CooRecorder) and have been adopted widely, none of these tools is completely open-source and free, slowing down further community-driven development. While ROXAS (von Arx & Carrer, 2014) provides a very powerful interface it depends on expensive libraries and is programmed in a discontinued programming language. CooRecorder provides a cheaper, albeit not free, and well-supported alternative to measure basic wood features, but does not provide its code base, prohibiting further development by the community. The only freeware, that we are aware of (e.g., ImageJ plug-ins), have not been widely adopted, presumably due to issues related to functionality, ease of installation and use, and/or support. As a result, a large part of the community either uses proprietary commercial software (e.g., WinDendro, CooRecorder) and many scientists still prefer manual measurements using “linear tables”, although the accuracy of digital analysis was already comparable to manual measurements about a decade ago for some species (Maxwell et al., 2011). Digital technology has advanced rapidly since and offers generally recognised benefits of simplified evaluation, corrections and amendment of previous measurements. Moreover, R packages such as *dlpR* (Bunn, 2008; Bunn et al., 2021), which allows tree-ring analyses from standard file formats, have enjoyed enormous success, arguably in part because the community is familiar with the R programming environment.

To move towards unlocking the wealth of underlying visual data from images of wood, we created an open source pipeline, the Wood Image Analysis and Dataset (WIAD), that integrates digital analysis and data archiving, while also laying the foundation for open-access sharing of images and derived data. WIAD is an easily-installed, simple R shiny application, whose graphical user interface can currently be accessed at https://wiad.science or run locally (see Code and Data availability for installation instructions). WIAD provides an interactive user-friendly interface for simple pre- and post-processing of wood images and metadata entry to facilitate a standardized workflow for labeling, measuring and detrending of radial growth. Both the web-interface and locally running copies allow making, evaluating and editing measurements and associate a minimal set of metadata (Table 1). When running WIAD locally, outputs are saved on the local machine enabling the upload and sharing of local collections at any point thereafter (e.g., upon acceptance for publication). Thus, the user can embargo the data by running a copy locally and only uploading the data upon publication as per terms of use. When using the web interface, data is archived straight away to the WIAD repository.

**Table 1:** The following metadata is associated and archived with each image and derived data on the Wood Image Analysis and Database (WIAD). Metadata can be uploaded using a metadata form or filled in manually. Data types can be character strings (C), dates (D), integers (I), logical variables (L), or doubles (N). The units column displays the assumed units for each metadata field. The optional column specifies whether the metadata is optional (✓), required (*) or provided by the software (✗).

| Variable name | Type | Units | Optional | Description |
| --- | --- | --- | --- | --- |
| Version | C | NA | ✗ | Version of WIAD used to produce this file. |
| Image file name | C | NA | ✗ | Name of the image file |
| Name | C | NA | ✓ | User's full name |
| Email address | C | NA | ✓ | Email address of user |
| Species | C | Unitless | ✓ | Genus or, if available, species name after |
| Sample date | D | yyyy-mm-dd | ✓ | Date of sample collection |
| Growing season | C | NA | * | Factor determining whether the growing season had not started (default), only started, or already finished |
| Schulman Shift | L | NA | * | Determine whether a shift of the associated year (mainly necessary in Southern Hemisphere) should be performed according to Schulman (1956). The default is no shift. |
| Scan resolution | I | Dots per inch | ✓ | True (not interpolated) resolution of the scan |
| Site location | C | NA | ✓ | Detailed description of site location including area, town, and country |
| Latitude | N | decimal ° | ✓ | Bounding latitudes in decimal notation for the sample location |
| Longitude | N | decimal ° | ✓ | Bounding longitudes in decimal notation for the sample location |
| Site ID | C | Unitless | ✓ | Unique identifier used by contributor for the specific site |
| Plot ID | C | Unitless | ✓ | Unique identifier used by contributor for the specific plot |
| Sample ID | C | Unitless | ✓ | Unique identifier used by contributor for the sample |
| Sample height | N | m | ✓ | Height aboveground at which the sample was taken |
| Sample azimuth | I | ° | ✓ | Azimuth angle at which the sample was taken |
| Sample note | C | NA | ✓ | Any additional notes pertaining to the specific sample |
| Collection | C | NA | ✓ | Name of the collection to which this sample belongs |
| Contributor | C | NA | ✓ | Full name of the contributor who led the collection effort and owns the collection |
| Pith in image | L | NA | * | Determine whether the image and data series contains the pith. By default it is assumed that there is no pith in the image. |
| Bark first | L | NA | * | Determine whether the measurement series starts at the bark (default) or at the pith |

In the following sections, we explain WIAD’s current functionality, provide an illustrative example of how it can be used, describe anticipated developments and the future potential of such a community-developed platform. Community-developed platforms to archive, share, and analyse images in a transparent and reproducible way is applicable for a vast array of visual data in ecology, such as leaf or root scans, and have proven very successful for digital images of canopies (Hufkens et al., 2018; Richardson et al., 2018). WIAD aims to provide a blueprint of an integrated repository and analysis framework to advance the ecological data revolution using agile software development and state-of-the-art practices in scientific software development (Ahmed et al., 2014; List et al., 2017; Wilson et al., 2014, 2017).

## 2. WIAD functionality

After the initial installation as an R package (available on CRAN and github, see Code and Data availability for installation instructions) including its dependencies (most prominent R and the shiny package), WIAD can be run locally by one simple command. Alternatively, WIAD can be accessed through a web-interface at https://wiad.science. The application currently has four tabs: Toolbox, Plot Board, About and Fair use policy.

### 2.1 Toolbox

The toolbox provides basic interactive image manipulation tools (Fig. 1), which require the upload of an image in TIFF, PNG or JPEG format. Once uploaded, the image can be rotated, processed with colour transformations (i.e., display true colour, total brightness, or blue channel only), and enlarged using a zoom bar (digital magnification up to a maximum image width of 20 000 pixels). The user can enter - either manually or via the upload of an XLSX, JSON or CSV file - and needs to confirm the metadata, which is then associated with the image and all derived data. To convert the distances in the image from pixels to micrometers, the user needs to provide the true scan resolution (in dots per inch), otherwise distances will be provided in pixels.

**Figure 1.**
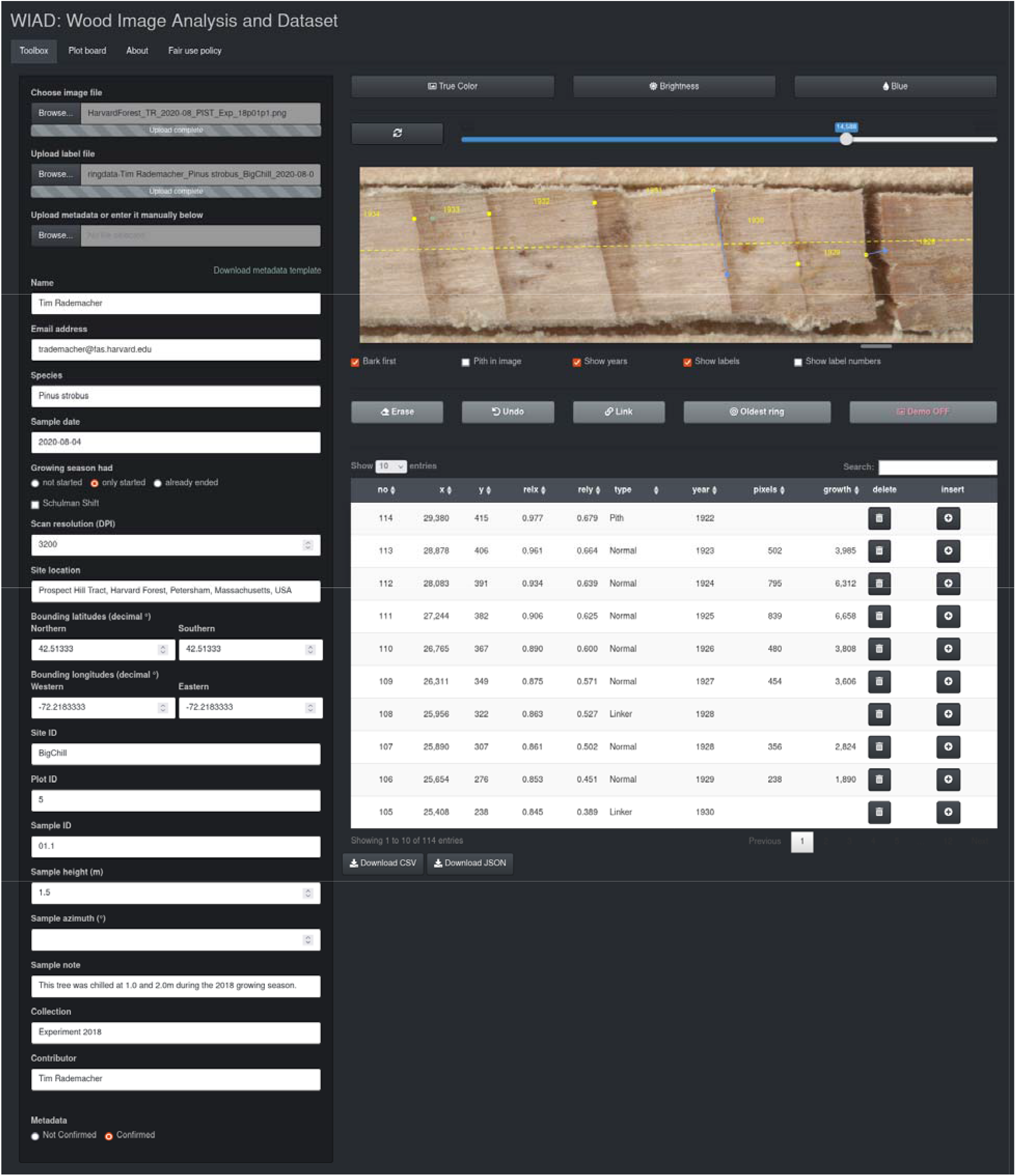
Screenshot of WIAD Toolbox with an example image of white pine and uploaded labels from a previous measurement. The metadata automatically populates the fields in the left column, but needs to be confirmed with the radio button below. Normal labels are overlaid onto the image in yellow, miscellaneous labels in turquoise and link labels in blue.

**Figure 2.**
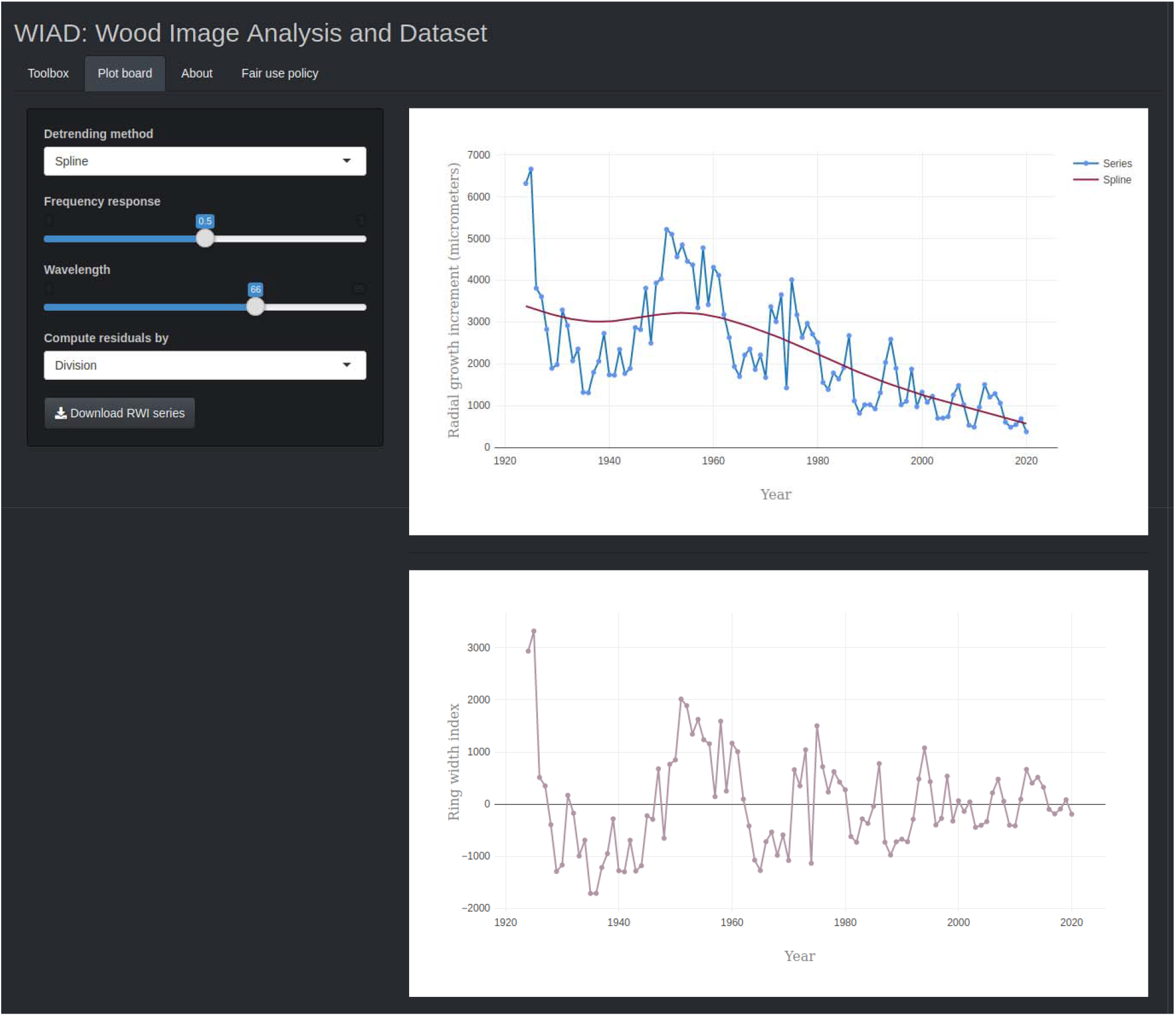
Screenshot of the WIAD Plot board showing the controls for detrending on the left and interactive plots of the radial growth increments (blue line and markers; top graph) including a detrending curve (red curve; top graph) and ring-width indices (mauve line and markers; bottom graph).

Once the metadata is confirmed, the toolbox allows setting, deleting, and inserting five types of labels through a simple graphical user interface. A simple click on the image sets a normal label marking the end of a growth increment (overlaid in yellow), a double click sets a miscellaneous label (overlaid in turquoise), which marks miscellaneous features that can be selected from a pop-up dialog. Radial growth (i.e., one-dimensional distance between labels in micrometers) and association of a particular year with each annual growth increment (including a one-year shift when desired in the southern hemisphere sensu Schulman (1956)) are performed automatically and displayed in a data table below the image. The user can delete a specific label, the last label, or all labels with the “Bin” of the specific data table row, the “Undo”, or the “Erase” button below the image, respectively. The user can switch a normal label to a link label (overlaid in blue) by pressing the “Link” button below the image, once the label was set. Links can bridge gaps in the sample, when two consecutive linker labels are used, or relocate the start point for a measurement (as visible in Fig. 1). Another action button converts a normal label to mark the oldest completely visible ring or pith (overlaid in red). Checkboxes below the image represent a choice whether the labelling started at the bark side of a core, whether the image contains the pith, and whether years, labels and/or label numbers should be displayed. Additional labels, including missing rings, can be inserted before any existing label by clicking on the “Add” button of the respective row of the data table. Users can also upload images, metadata and labels to evaluate previously analysed images or add images that were analysed offline to the online archive. Finally, users can download a CSV or JSON file, which contains labels, derived measurements and metadata (the latter is not included in CSV format) to save their progress or for further analysis offline.

To use the tool for teaching or demonstration, the “Demo” button switches demonstration mode on/off. In demonstration mode, an example image is automatically loaded, but can be replaced, and data or images are neither archived locally, nor in the online database.

### 2.2 Plot board

The WIAD plot board tab provides two interactive plots to quickly assess the derived growth data and detrended ring-width indices. The upper plot displays the measured annual radial growth increments over time, while the second graph shows ring-width indices based on residuals from an expected growth curve (Cook & Peters, 2016). The user can select a detrending curve from the *dplR* package (Bunn, 2008; Bunn et al., 2021) to obtain ring-width indices via controls on the left and examine them in real-time. Finally, a JSON file including the labels, derived measurements, detrended ring-width indices, detrending parameters, and metadata can be downloaded.

### 2.3 About and Fair use policy

The “About” tab provides basic information about WIAD, whereas the “Fair use policy” tab outlines the conditions of use with regard to the tools and data. WIAD code base is publicly available for noncommercial reuse under the Affero General Public License v3 license (https://www.gnu.org/licenses/agpl-3.0). Accumulated images, derived data, and metadata will be made publicly available every couple of years after thorough review. We request that the WIAD core development team and data contributors are appropriately acknowledged in work using WIAD by citing this paper and the appropriate software citation. Contributors’ names and email addresses are attached to each datum in the repository through the accompanying metadata (if they were provided) and we expect data users to contact data contributors. Data contributors will have a deeper understanding of the data and additional insight; thus we consider them crucial collaborators for any data-user.

Despite higher resolution scans being generally desirable, as they capture more details, which may lead to additional uses in the future, larger images also require more storage. Image rendering is the computational bottleneck of WIAD, hence larger images also slow down processing. Therefore, we recommend a scan resolution of 3200 dpi, which balances the tradeoff between visual quality and file size for standard increment cores. To reduce storage requirements and optimise processing speed images should always be cropped (using open-source image manipulation programs like GNU Image Manipulation Program) to only the region of interest prior to upload. For particularly large samples (e.g., cross-sections), hence files, segmentation into multiple images can be considered for faster processing and reduced storage. When pre-processing scans, care must be taken to avoid changing the image resolution or features. Due to storage limitations, maximum image size of the web-interface is currently limited to 30 MB (i.e., sufficient for a 3200 dpi scan of a 30 cm long and 6mm wide sample). When running WIAD offline, the maximum image size can be adjusted via the maxImageSize variable in the *server*.*R* file.

## 3. Example: Measuring ring width and density fluctuations in white pine

To provide an example data set we measured and uploaded images for ten mature white pines (*Pinus strobus* L.) from Harvard Forest, USA. For this data set, we collected two increment cores at breast height using a standard increment corer (5.15mm, Hagl□f Company Group, Långsele, Sweden). The cores were glued to wooden mounts and sanded with progressively finer sandpaper (from 80 to 8000 grit) until visual features of the wood were clearly visible by the naked eye. A subset of these images and derived data of 250 MB can be downloaded with the getExampleData function. To download the entire example dataset (1.83 GB) the function’s “dataset” parameter must be set to “full”.

### 3.2 Imaging samples

Samples were scanned with an Epson Perfection V500 Photo Scanner (Long Beach, California, USA) at 3200 dpi with an advanced reflective colour-calibration target (10×15cm target printed on Kodak Professional Endura, LaserSoft Imaging, Kiel, Germany). The colour calibration target allows for tracking and potential future correction of drift in colours over time and between scanners. We recommend digitising samples on a flatbed scanner with a charge-coupled devices (CCD) imaging array. The CCD imaging arrays have a larger depth of focus compared to the increasingly popular complementary metal oxide semiconductor arrays, although the latter may outperform CCD arrays in the future (Janesick & Putnam, 2003).

## 4. Anticipated developments and future potential

WIAD complements existing tools by providing a free and open-source alternative in a widely used programming language, which facilitates further community-driven development. While size limitation of the online versions (i.e., 30 MB) and the basic nature of the processing and analysis tools in the current implementation are clear limitations, we plan to extend the framework to provide a pipeline that allows users to process, measure, and analyse images as well as archive the raw, derived, and meta-data all in one place. Such a pipeline will improve transparency and reproducibility of studies based on wood images by enabling easy sharing of data and provenance tracing. We intend to regularly publish curated data sets from the WIAD archive every couple of years. This approach has proven a workable solution for the PhenoCam network (Richardson et al., 2018; Seyednasrollah et al., 2019), as it reduces the need for intensive and immediate quality control and provides cleaned and citable versions of data sets to the community. In the hope that other developers will join our efforts, we outline a few specific directions below.

### 4.1 Exploiting the potential of already existing collections by digitisation

Wood is arguably the most abundant biological tissue on earth (Bar-On et al., 2018) and records information about past climates (Mann et al., 2008), disturbances (Bergeron et al., 2002; Lorimer, 1984) and biotic interactions (Cook, 1987). Scientists of multiple disciplines (ecology, physiology, archeology, etc.) have collected millions of wood samples in the form of increment cores, micro-cores (Rossi et al., 2006), and stem cross-sections, which are archived in laboratories across the world. Publicly available digital images of collected samples would move these fields to a big data domain and provide the foundation for major advances in our understanding of the process of wood formation, its abiotic and biotic constraints, and resulting anatomy. Improved archives will be instrumental for teaching the next generation of scientists, reconstructing past environments and will improve forecasts of future carbon sequestration that are based in ecological realities. Digitisation and data sharing have multiple proven benefits such as accelerating science and improving trust in science (Michener, 2015; and references therein). WIAD paves the way to digitise large collections of wood samples and contribute to improved archives. Additionally, WIAD can be used as a teaching tool in demonstration mode, especially if users share specific image libraries addressing teaching goals (e.g., containing drought-induced intra-annual density fluctuations).

While WIAD provides the foundational tools, several aspects would profit from refinement such as more comprehensive metadata, association with ancillary data, batch processing of images and integration of data provenance. WIAD associates images with a minimal set of metadata (Table 1) and we plan to integrate handling of more comprehensive metadata using the Ecological Metadata Language (Fegraus et al., 2005; Jones et al., 2006). Similarly, WIAD intends to comply with Darwin Core Standard (Wieczorek et al., 2012) to ensure compatibility with other data collections of biological diversity. Finally, future versions should allow the upload of ancillary data such as data of non-visual chemical or physical properties (isotopic composition, nonstructural carbon concentrations, lignin content, etc.).

### 4.2 Streamline data processing through automation and enable community science

To further automate the processing of digital analysis of wood samples, low hanging fruit will be the automatic detection of visual properties. Initial algorithms correctly identified almost 100% of rings in four conifer species, roughly 85% in six diffuse-porous species, but only 40-50% in two ring-porous species (Fabijańska et al., 2017; Subah, Derminder, & Sanjeev, 2017), despite being based on comparatively small and homogenous training data. Recently, convolutional neural networks have also succeeded at segmenting anatomical sections of wood (Garcia-Pedrero et al., 2018). The integration of visual computing algorithms into WIAD to automate the analysis and reduce analysis time, by suggesting ring boundaries to the user after image upload, is under development. For this purpose, we are currently creating the first fully labeled image library of sufficient size (more than 100 000 labels in 1000 images) to train a robust ring detection algorithm that works for ten species including ring-, diffuse- and non-porous wood. Specifically, we adopt the U-Net model (Ronneberger et al., 2015) to predict the pixel-wise probability of ring boundaries. Since models trained with the default pixel-wise cross-entropy loss usually yield boundaries with blurry artifacts, we incorporate a perceptual discriminator energy function to improve the sharpness of the boundaries while preserving the connectivity, following the high-level idea of Mosinska et al. (2018). Beyond identifying rings, the identification of other visual features such as wood morphology (e.g., ring-porous, diffuse-porous, and non-porous), density fluctuations or frost and fire scars, could equally be automated using similar techniques. Another useful addition would be the integration of cross-dating tools that help to visually and statistically crossdate multiple measurement series such as xDateR (Bunn, 2008, 2010) or ringdater (Reynolds et al., 2020). Similar to suggestions of ring boundaries, fast fuzzy subsequence matching (Gong et al., 2019) and unsupervised anomaly detection algorithms for either time-series (Ren et al., 2019) or high dimensional data like raw images (Wang et al., 2019) could be used to assist in the crossdating and identification of missing rings, which would further increase processing speed and improve reproducibility of results.

Beyond streamlining sample processing, these features could enable scientists and potentially even citizen scientists to contribute to the repository in the long-term. However, sufficient skills of the software must be demonstrated rigorously before expert opinion can be meaningfully complemented. Nonetheless, the general public could already help in the labelling of datasets and thereby provide training data for the refinement of algorithms. Crowd-sourcing through a community science platform (e.g., Zooniverse) could be used to label training data and evaluate annotations of existing collections. Community-science projects come with tradeoffs in data quality, privacy protection, resource security, transparency, and trust by citizens (Anhalt-Depies et al., 2019), which we will consider during further development.

In conclusion, WIAD provides simple tools and a framework for further development to make analysis of wood images more transparent and reproducible. By offering integrated data processing and archival services, platforms such as WIAD facilitate analysis by making data accessible, tracking provenance, assuring metadata standards and, therefore, creating data products that can be used, amalgamated, scaled, built upon, and modified seamlessly.

## Author contributions

TR conceived the idea and developed it with BS and DB. BS developed the code with input and additions from TR and DB. TR cored the trees, mounted and sanded the cores. EM, TM, and TR scanned images and tested the tools. JC, ZL, and DW developed the algorithm for automatic boundary detection. All authors provided ideas on how to improve the tools. TR led the writing of the manuscript. All authors contributed critically to drafts and approved the final manuscript.

## Code and data availability

All data is publicly available through getWIADExampleData function built into the package. The R package requires R to be installed and underlying code are also publicly accessible under a AGPL v3 license on github at https://github.com/bnasr/wiad and is easily installed via CRAN:

> utils::install.packages(‘wiad’,repos=‘https://cran.us.r-project.org’)

Or directly from github using:

> if (!require (devtools)) {install.packages (‘devtools’)}
>
> devtools::install_github (‘bnasr/wiad’)

The interactive mode can be launched from an interactive R environment by one of the following commands.

> Library (wiad)
>
> Launch ()

Or

> wiad::Launch()

or form the command line (e.g., shell in Linux, Terminal in macOS and Command Prompt in Windows machines) where an R engine is already installed by:

> Rscript -e “wiad::Launch(Interactive = TRUE)”

Calling the Launch function opens up the WIAD app in the system’s default web browser.

## Acknowledgements

TR and TM acknowledge the support from Microsoft “AI for Earth” grant program (271089). AR, EM and TR acknowledge support from the National Science Foundation (DEB-1741585, HF LTER DEB-1237491 and DEB-1832210). DB acknowledges support through the Swiss National Science Foundation (PSBSP3-168701) and the Harvard Forest Bullard Fellowship. EM was also benefited from NSF DBI-1459519 and the Harvard Forest Summer Research Program in Ecology. We also thank Patrick Baker, Greg Caporaso, Allyson Carroll, Patrick Fonti, George Koch, Rubén D. Manzanedo, Steve Sillett, Laura G. Smith, Yiping Zhang for their valuable feedback.

